# Predicting Phenotypes From Novel Genomic Markers Using Deep Learning

**DOI:** 10.1101/2022.09.21.508954

**Authors:** Shivani Sehrawat, Keyhan Najafian, Lingling Jin

## Abstract

Genomic selection models use Single Nucleotide Polymorphism (SNP) markers to predict phenotypes. However, these predictive models face challenges due to the high dimensionality of genome-wide SNP marker data. Thanks to recent breakthroughs in DNA sequencing and decreased sequencing cost, the study of novel genomic variants such as Structural Variations (SVs) and Transposable Elements (TEs) become increasingly prevalent. In this paper, we develop a deep convolutional neural network model, `NovGMDeep`, to predict phenotypes using SVs and TEs markers for genomic selection. The proposed model is trained and tested on samples of *A. thaliana* and *O. sativa* using *k*-fold cross-validation. The prediction accuracy is evaluated using Pearson’s Correlation Coefficient (PCC), Mean Absolute Error (MAE), and Standard Deviation (SD) of MAE. The predicted results showed higher correlation when the model is trained with SVs and TEs than with SNPs. `NovGMDeep` also has higher prediction accuracy when comparing with conventional statistical models. This work sheds light on the unrecognized function of SVs and TEs in genotype-to-phenotype associations, as well as their extensive significance and value in crop development.

## Introduction

The genotype of an organism is described by its entire genetic composition. Genotype can also refer to the set of alleles contained within the genome. The phenotype, or observable characteristics, are influenced by the genotype’s expression in the cell. Generally, three variables influence a phenotype: the most effective are genes; the inherited epigenetic factors and acquired environmental factors are the other two [34].

Because of the success of Genome-Wide Association Studies (GWAS), there is a growing interest in using genotype data to predict complex traits like diseases. According to some recent GWAS analyses only a few common SNPs are involved in most diseases and these associated SNPs can only explain a small percentage of the disease susceptibility [19]. Early plant genomic investigations were hampered by technological limitations and a lack of high-quality reference genome assemblies, which prohibited a detailed analysis of both SVs and TEs in plants. Recent improvements in genomic technology, notably long-read sequencing, offer the generation of high-quality plant genome and pangenome assemblies, as well as exposure to a diverse set of SVs for evaluating their possible significance in plant phenotypic diversity [38]. Similarly, the transposition of TEs, which are common in most eukaryote genomes, accounts for a considerable amount of genomic variation. As a result, TE-derived molecular markers are useful resources for unraveling genomic variations in both plant and animal [28]. This work aims to shed light on the unrecognized function of SVs and TEs in genotype-to-phenotype associations, as well as their extensive significance and value in crop development.

Predicting crop phenotypes is an effective step in explaining crop behavior using insights from genome-wide markers. Genomic selection (GS) is a potent method to enhance quantitative traits since it uses genomic-estimated breeding values of individuals generated from genome-wide markers to select candidates for the next breeding cycle [21].

The importance of predicting complex traits using large volumes of genomic data has led to the development of novel machine learning models. The conventional genomic prediction models are rrBLUP (Ridge Regression Best Linear Unbiased) [8] and gBLUP (Genomic Best Linear Unbiased Prediction) [6]. However, these conventional predictors typically assume that the genotype random effects follow a normal distribution, and each genotype’s contribution to phenotypes is considered as an independent attribute. It is, however, unknown in practice how genotype effects behave and they may not follow a strict distribution. Moreover, these conventional statistical models do not consider non-linearity between the variables, as SNPs may also interact with other SNPs to cause complex diseases or traits as a result of epistasis. While these models face challenges due to the high dimensionality of genome-wide marker data and interactions between alleles, GS can benefit from Deep Learning (DL), which provides novel approaches to process noisy data [18] and handle nonlinearity [15].

For DL models, studies have shown that these methods base their predictions on overall genetic relatedness, rather than on the effects of particular markers [32]. Moreover, DL models offer advanced feature engineering and extraction capabilities [41], and therefore have the potential to find hidden patterns in large-scale datasets [22]. As a simplest DL model, in a multi-layer convolutional neural network, each layer consisting of many neurons, represents complicated relationships among the large datasets. The different neurons in each layer, receive information from the lower hierarchical layers, which is triggered by predefined activation functions. Activation functions bring non-linearity to the output of each layer to determine the input of the next layer [23]. Deep learning-based models has shown promising results in wide variety of tasks such as object detection [40], speech recognition [7], natural language processing [37], and etc.

### Existing Methods in Genomic Selection

Broadly speaking, GS methods can be classified into conventional statistical models and deep learning methods. In this section, we briefly summarize a few models in each category. These models are compared with our proposed `NovGMDeep` model in Section 3 to demonstrate the prediction of phenotypes using SV, TE, and SNP markers.

One of the most widely used statistical models for GS is rrBLUP [8]. The model uses the linear regression algorithm which takes the genotype matrix as the input and predicts the phenotype vector. The ridge regression technique is used to estimate the effects of all genotypic markers, and these effects follow a normal distribution with a non-zero variance.

The gBLUP model [6] is another statistical model for estimating an individual’s genetic performance based on its genomic associations. A genomic relationship matrix is used in gBLUP to represent genomic markers. To make predictions, the genotype matrix specifies covariance between individuals based on observed similarity at the genomic level.

The DeepGS model is developed using a deep Convolutional Neural Network (CNN) with architecture including a 1D-convolutional layer, a 1D-max-pooling layer, combination of a few dropouts and fully connected layers [20]. The final output represents the predicted phenotypic values for the analyzed individual markers. The DeepGS model is trained using 10-fold cross-validation. They used the backpropagation algorithm [29] with the learning rate of 0.01, the momentum of 0.5, and the weight decay set to 0.00001 to optimize the parameters of the model. They trained the model for 6000 epochs. The DeepGS model utilizes hidden variables to collectively represent features in the genome-wide markers while making predictions.

The quantitative phenotypic prediction problem was treated by G2PDeep as a regression problem [39]. Data on zygosity and SNPs are fed into the model. The model is made up of a dual-CNN layer and a fully connected neural network. The encoded genotypes are transferred by dual-CNN, which has two concurrent series of CNN layers with the kernel sizes of 4 and 20, followed by another CNN layer of size 4 to enhance marker representation. Both CNN streams are aggregated and fed into the following CNN layer to complete the feature extractor section of the model. In the end, fully connected layers with 512 and 1 neurons serve as regression blocks for predicting phenotypes in the model.

### Genomic Selection using Novel Genomic Markers

The above-discussed methods only consider the SNP variants as genomic markers in predicting the phenotypes. For standard linear models, the number of genome-wide SNP features largely exceeds the sample size, leading to overfitting. To mitigate these limitations, instead of SNPs, we utilize SVs and TEs to represent genomic variations in two separate experiments and explore their effectiveness in genomic selection. We have provided an overview of SVs and TEs in Section 1 and explained their importance in phenotypic prediction in plants.

Our study illustrates the need to look beyond SNPs to understand evolutionary processes and how SVs and TEs can help us understand variation within species or during early divergence. The genotype-to-phenotype predictions identified using SVs and TEs will be useful to investigate *A. thaliana* and *O. sativa* evolution and trait architecture. We develop `NovGMDeep`, a 1D deep convolutional neural network, to predict the different phenotypes from novel genomic markers-SVs and TEs. Unlike the statistical models, our model learns the complex relationships between genome-wide markers and phenotypic traits from the training data. With the advanced deep learning technology, the model evades the overfitting of the data using the convolutional, pooling, and dropout layers hence decreasing the complexity of dimensional genomic markers.

This paper is organized as follows. In the Material and Methods section, first, we discuss our data sources, second, we present our proposed DL model, and third, we present the data representation for the proposed model in detail and its architecture and optimization strategies. Then, in the Results section, we discuss the prospective results by comparing the results with all of the mentioned statistical and DL models. Finally, we draw some conclusions and discuss how predicting phenotypes using SVs and TEs has more impact than using SNPs in the Conclusion section.

## Materials and Methods

### Data Sources

#### *Arabidopsis thaliana* dataset with SV markers

The flowering plant, *A. thaliana*, is an ideal plant species for researching genotype-phenotype-environment interactions because their naturally inbred strains allow for repeated phenotyping and have adapted genotypes under a variety of controlled circumstances. *A. thaliana* is quite appealing in scientific research for surveying the molecular and phenotypic effects at the species level [33]. It is found in a wide variety of environments around the world, and representative accessions have been extensively phenotypically and genetically characterized. The 1001 Genomes “A Catalog of *A. thaliana* Genetic Variation” [27] is a database where whole-genome sequencing has been performed on over 1000 *A. thaliana* accessions which makes the genomic selection and genome-wide association studies in this species possible.

The full VCF variant files containing the structural variants data for the 1, 301 *A. thaliana* samples are publicly available on European Variation Archive (PRJEB38975) [10]. They were used in this study, as well as the SNPs and indels for 1, 135 accessions of *A. thaliana* [3]. The SV dataset includes several types of SVs, such as deletions, duplications, and inversions. The lengths of these variations span from 50 bp to 10 kb.

The phenotypes of matched samples were downloaded from AraPheno, the 1001 genomes project [31]. Flowering time, rosette leaf number, cauline axillary branch number, and stem length were used with their definitions listed as follows.

1. Flowering time: Generally scored as days from seeds in the soil until the first open flower.
2. Rosette leaf number: A shoot system morphology trait which is the number of leaves in the shoot system of *A. thaliana*.
3. Cauline axillary branch number: A shoot axis morphology trait which is the amount or pattern of branches arising from the shoot axis.
4. Stem length: A stem morphology trait often measured from the soil surface to the highest point on the stem.

#### *Oryza sativa subsp. japonica* dataset with TE markers

*O. sativa*, rice is a monocotyledonous flowering plant in the *Poaceae* family that is one of the world’s most significant agricultural plants, providing the primary source of nutrition for half of the world’s population. *Oryza sativa subsp. japonica* is one of three primary rice subspecies: *indica, javanica*, and *japonica*. The grains of *O. sativa* are short and high in amylopectin, causing them to stick together when cooked.

The TE markers computed from the 176 *O. sativa* accessions were used in this study [35]. The SNPs for the same number of accessions were also obtained in this study in comparison to TE markers.

Four phenotypes are used in the study: panicle length, spikelet number, days to heading, and plant height measured for the same accessions [36].

1. Panicle length: The length measured from a terminal component of the rice tiller which is an inflorescence called a panicle.
2. Spikelet number: The number of panicle rice spikelets, which develop into grains.
3. Days to heading: Characterized together by the vegetative growth phase. It is the period from germination to panicle initiation and the reproductive phase of rice development, meaning the time from panicle initiation to heading.
4. Plant height: Defined as the shortest distance between the upper boundary (the highest point) of the main photosynthetic tissues (excluding inflorescences) and the ground level.

`NovGMDeep`: a Deep Learning Model for Genomic Selection using Novel Genomic Markers

We present, `NovGMDeep`, a deep one-dimensional (1D) convolutional neural network to predict phenotypes in this study. The model is trained separately with three different types of data: SVs, TEs, and SNPs. We evaluate the model by calculating Pearson’s coefficient of correlation among the predicted and observed phenotype values. The results show that SVs and TEs are more informative for phenotype prediction than SNPs.

### Genomic Data Representation

Genotypes in plant and animal species are represented as either haploid or diploid. A *haploid* genotype comprises a single set of chromosomes for generating the phenotype whereas a diploid genotype contains two sets of chromosomes, each of which is required for phenotype generation. Genotypes are usually represented as ‘0|0’ or ‘0*/*0’, where ‘|’ and ‘*/*’ indicates the phased and unphased genotypes respectively. To gain a complete picture of genetic variation, phasing entails separating maternally and paternally inherited copies of each chromosome into haplotypes. A genotype with at least one set of explanatory haplotypes is referred to as a phased genotype. Unphased genotypes are those for which no set of explaining haplotypes have been identified [24]. The genotype is considered to be *homozygous* if all the haplotypes for a specific site have the same value. Otherwise, the genotype is considered *heterozygous*.

#### SV marker representation

*A. thaliana* is a diploid species and has phased genotypes as ‘0|0’ and ‘1|1’ which means homozygous for reference allele and homozygous for alternate allele respectively. Although SVs are smaller in numbers than SNPs, which account for almost a million variant sites per genome, they are more informative, as they account for a higher number of nucleotide differences due to their size. Different types of SVs such as inversions, duplications, and deletions can capture the genomic variations better than the SNPs. We propose a strategy as shown in formula 1 to represent the SVs. The SV genotypic data for inversions, duplications, and deletions is transformed using one-hot encoding as binary arrays. In one-hot encoding, given a dataset with different features, the encoder finds the unique values per feature and transforms the data to a binary one-hot vector [26]. Different arrays for different types of SVs were chosen so that the details of genomic variations were well represented and understandable by the model. For the SNP and indel data, the same one-hot encoding strategy was used, 0|0 for reference allele, 1|1 for alternate allele, and .|. indicating the missing genotypes. As shown below, INV represents inversions, DUP represents duplications, and DEL represents deletions:

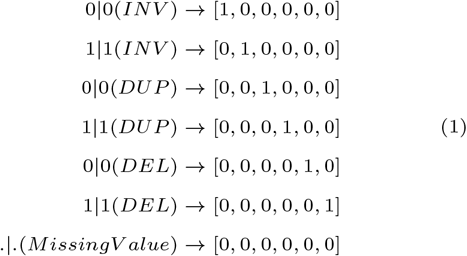

For the SV dataset, the data is represented in the format of a 3D matrix *A*(*x × y × z*) where *x* represents the accessions, *y* represents the SV/SNP markers, and *z* represents the size of one-hot-encoded arrays. As one example, in the *A. thaliana* SV dataset, there are 914 accessions, 155, 440 genotypes, and 6 variant types as defined in formula (1), therefore, *x* = 914, *y* = 155, 440, and *z* = 6 respectively. As another example, for the *A. thaliana* SNP dataset, *x* = 923, *y* = 500, 000 (only 500, 000 SNP genotypes were selected randomly from the 10 million genotypes to save computational resources), and *z* = 2 respectively.

#### TE marker representation

*Oryza sativa* is a diploid species and has unphased genotypes. The TE and SNP markers in the raw dataset were coded as integers 1, 0, and -1 which means the reference genotype (1*/*1), the first (most common) alternative genotype (0*/*1), and the second most frequent alternative genotype (0*/*0) respectively. A similar strategy is used to encode TE and SNP variants using one-hot-encoded binary arrays as shown in formula (2).

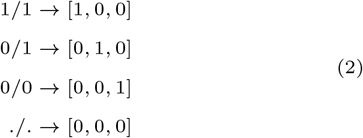

For the TE markers, the data is represented in the similar format of a 3D matrix *B*(*u × v × w*) where *u* refers to the accessions, *v* indicates the TE/SNP markers, and *w* shows the depth of one-hot-encoded vector. So, for the *O. sativa* TE dataset *u* = 176, *v* = 6, 074, and *w* = 3 respectively. And for the *O. sativa* SNP dataset, *u* = 176, *v* = 493, 882, and *w* = 3 respectively.

### Model Architecture

The proposed deep CNN model has four 1D convolutional layers, a single 1D max-pooling layer, a flatten layer and one dropout layer followed by a fully connected layer (Figure 1).

**Fig. 1.**
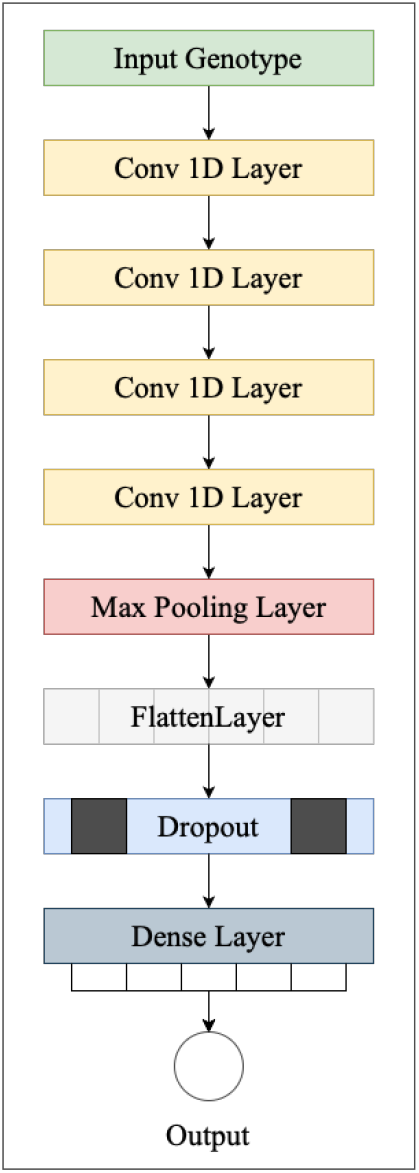
`NovGMDeep` architecture. The first convolutional layer has 16 filters of kernel size five. The second layer comes with 32 filters of kernel size five. The third one has 64 filters of kernel size three. And the last convolutional layer has 128 filters of kernel size three. The max-pooling layer has a pool size of two and a stride of two. At last, there is one fully connected layer as the regressor with one output neuron.

As the backbone of the model, we use four 1D convolutional layers of the following sizes. The first layer includes 16 filters of size five. For the next layers, we double the number of filters in each layer and reduce the filter sizes of the last two layers to three. If the input to the convolution layer is represented as *X*, filter as *F*, bias as *B*, and the output of the convolution layer as *Y*, the convolution operation between *X* and *F* including bias is defined in equation (3).

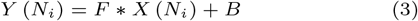

where *N* is the batch size and *∗* is the cross-correlation [5] operator. For all the convolutional layers, ReLU activation function [1] (Equation 4), LecunNormal kernel initializer [12], and L1L2 kernel regularizer [17] is used with same padding.

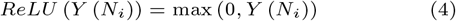

To reduce the high dimensionality of the prominent features of the last convolutional layers, a one-dimensional max-pooling layer is used after the convolutional layers with a window size of two and a stride of two. It minimizes the amount of the input to the following layer by taking only the maximum values as the most important features, which cut the size of the input into the following layer in half [2]. Then the extracted feature map from the model’s backbone is flattened using a flatten layer and fed into the fully connected layer which is our phenotype predictor. Fully connected layers link every node in the input to every node in the output, but they do not capture spatial information. Due to the high number of input features which results in a large-size feature map, the dropout layer with the drop ratio of 0.6 is used before the fully connected layer to reduce computational complexity and avoid overfitting. We use Adam [16] optimizer with a learning rate of 0.0003. The Adam optimization algorithm is applied to iteratively adjust the network weights based on training samples. Moreover, we utilized mean absolute error as the loss function. The `NovGMDeep` is developed with Keras [11], an open-source software library that provides a Python interface for artificial and deep neural networks. The source code of `NovGMDeep` is publicly available on Github (https://github.com/shivanisehrawat4/NovGMDeep.git).

The model predicts quantitative phenotype as a real number which is the model’s output. The predicted phenotypes are later compared with the ground-truth values to evaluate the model performance. As the evaluation metric, we use Pearson’s coefficient of correlation (PCC) between the predicted and observed phenotypes. Five assumptions must be met before we calculate the PCC between two variables [25]. As far as our datasets are concerned, all of the conditions are true and the detailed analysis can be found in Appendix. PCC [9], normally denoted by *ρ*, is a measure of the linear correlation between two variables (*X* and *Y*), whose values range from -1 to 1, with 1 indicating total positive correlation, 0 indicating no correlation, and -1 indicating a total negative correlation. It is defined as the covariance of the two variables divided by the product of their standard deviations.

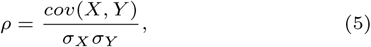

where *cov*(*X, Y*) is the covariance of *X* and *Y, σ*_*X*_ is the standard deviation of *X*, and *σ*_*Y*_ is the standard deviation of *Y*. PCC is a model-free method, and it, therefore, shows the nature of the data without depending on any of the existing models. It is commonly used in statistical analysis for the evaluation of the model and measuring the relationship between two variables.

## Results

We trained the `NovGMDeep` model on both SVs and SNPs variations of the accessions and their associated phenotypes for *A. thaliana*. We also trained the model on both TE and SNP data of *O. sativa*. We split all the samples into training and testing subsets with the ratios of 80% and 20% respectively. Note that we applied the 3-fold cross-validation [13] on the training sets of all the datasets in the model development step. The best models were saved while training for three rounds of cross-validation. Then, those best models were used to calculate the PCC value while testing. Later the average of the three PCC values obtained from testing was finally used in the results to evaluate the model.

PCC was used to find the correlation between the observed and predicted phenotypes. The correlation between the two quantitative variables assessed for the same samples is depicted by scatter plots in Figure 2 and Figure 3. The horizontal axis displays the true values of the phenotypes, while the vertical axis shows the values of the predicted phenotypes. Each value in the data is represented by a point on the graph. The strength of the association between the two variables is a key aspect of a scatterplot. The slope conveys information about the strength of the relationship between true and predicted values. When the slope is 1, the correlation between the two comparable variables is the strongest. Figure 2 shows the results between ground truth and predicted phenotypic values for `NovGMDeep` using SVs and SNPs respectively for predicting flowering time of *A. thaliana*. Figure 3 shows the results between observed and predicted phenotypic values for `NovGMDeep` using TEs and SNPs respectively for predicting heading time of *O. sativa*. These PCC graphs were created on the testing samples and show the accuracy of the proposed model in predicting phenotypes. The orange line in these graphs represents the trend of the relationship between the observed and predicted phenotypes (blue points) by the NovGMDeep model. This linear regression line shows the best fit between predicted and true values. As shown in Figures 2 and es 3, the predicted values which are close to the fitting line indicate a strong correlation, and values far from the fitting line indicate a weak correlation between the true and predicted values.

**Fig. 2.**
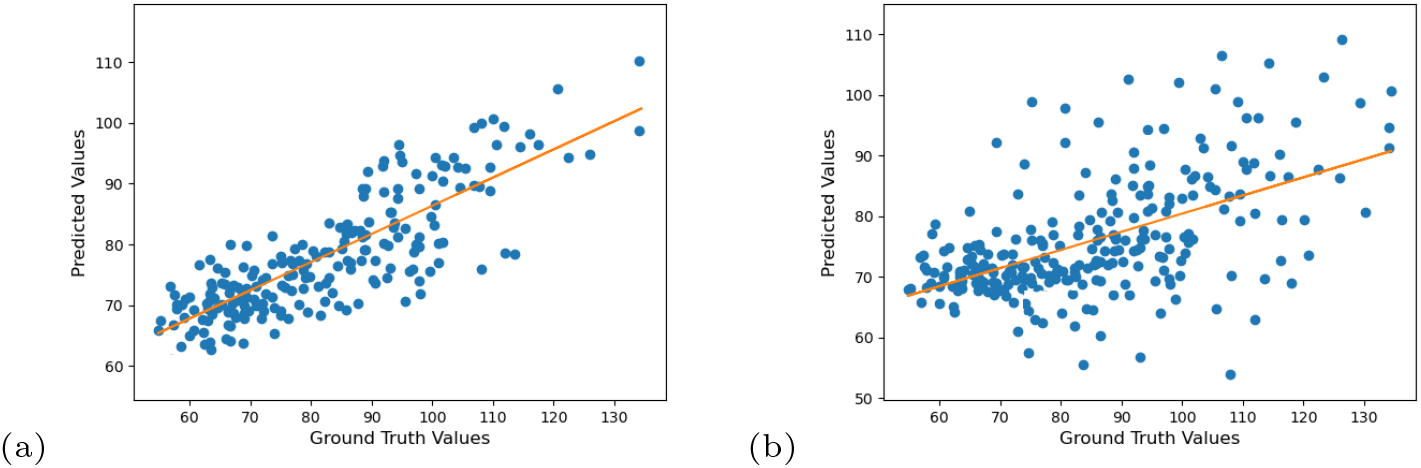
PCC analysis of *A. thaliana* for predicting flowering time using (a) SV and (b) SNP datasets. (a) The orange line represents the trend of the relationship between predicted and ground-truth values (blue points). The graph shows that predicted values by the model positively follow the ground truth values with a high confidence level (minimum fluctuation) in a positive direction. (b) The graph shows that predicted values by the model positively follow the ground truth values but with a low confidence level (more fluctuation).

**Fig. 3.**
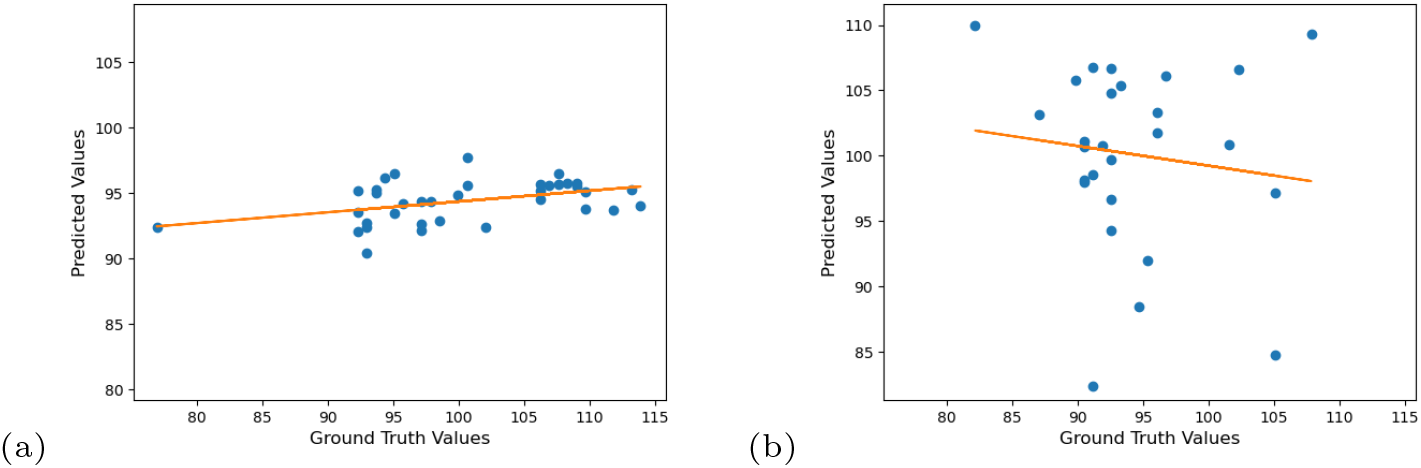
PCC analysis of *O. sativa* for predicting days to heading using (a) TE and (b) SNP datasets. (a) The graph shows that predicted values by the model are not following the ground truth values and have a very low confidence level. (b) The graph shows that predicted values by the model are negatively following the ground truth values and have no confidence level at all.

The aforementioned two statistical methods, rrBLUP and gBLUP, were evaluated with the same data to compare their overall prediction performance with `NovGMDeep`. As shown in Table 1, phenotypes predicted by `NovGMDeep` show the strongest correlations with observed phenotypes. Therefore, `NovGMDeep` outperforms the other statistical models in predicting phenotype with SVs data. Moreover, phenotypes predicted by `NovGMDeep` using SVs and TEs data show stronger correlations with observed phenotypes than the ones predicted using SNPs data. This shows the effectiveness of using SV and TE marker data instead of SNPs for phenotype prediction. Though, TE data showed the lowest PCC values compared to the two statistical models because of the mere 176 samples as there are not enough training samples for the DL model (Table 2).

**Table 1.** Pearson’s coefficients of correlations between the observed and predicted phenotypes on the testing set. SVs and SNPs are used to capture the genomic variations among the accessions of *A. thaliana*. Results show the average of the three PCC values taken from the three-fold cross-validation results of the model.

| Model | Flowering Time |  | Rosette Leaf Number |  | Branch Number |  | Stem Length |  |
| --- | --- | --- | --- | --- | --- | --- | --- | --- |
|  | PCC (SVs) | PCC (SNPs) | PCC (SVs) | PCC (SNPs) | PCC (SVs) | PCC (SNPs) | PCC (SVs) | PCC (SNPs) |
| *rrBLUP* | 0.664 | 0.520 | 0.653 | 0.547 | 0.675 | 0.513 | 0.690 | 0.502 |
| *gBLUP* | 0.657 | 0.516 | 0.662 | 0.512 | 0.683 | 0.530 | 0.681 | 0.514 |
| *NovGMDeep* | **0.788** | **0.594** | **0.742** | **0.563** | **0.761** | **0.581** | **0.759** | **0.572** |

**Table 2.** Pearson’s coefficients of correlations between the observed and predicted phenotypes on the testing set. TEs and SNPs are used to capture the genomic variations among the accessions of *O. sativa*. Results show the average of the three PCC values taken from the three-fold cross-validation results of the model.

| Model | Days to Heading |  | Plant Height |  | Spikelet Number |  | Panicle Length |  |
| --- | --- | --- | --- | --- | --- | --- | --- | --- |
|  | PCC (TEs) | PCC (SNPs) | PCC (TEs) | PCC (SNPs) | PCC (TEs) | PCC (SNPs) | PCC (TEs) | PCC (SNPs) |
| *rrBLUP* | 0.439 | 0.281 | 0.391 | 0.275 | 0.387 | 0.294 | 0.405 | 0.209 |
| *gBLUP* | 0.417 | 0.212 | 0.402 | 0.266 | 0.371 | 0.282 | 0.387 | 0.215 |
| *NovGMDeep* | **0.436** | **-0.014** | **0.406** | **0.002** | **0.372** | **-0.169** | **0.397** | **-0.148** |

The mean absolute error (MAE) metric is used to calculate the validation loss on the test set. MAE refers to the magnitude of difference between the prediction of observation and the true value of that observation.

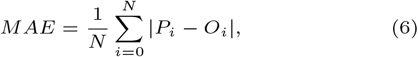

where *N* represents the total number of individuals present in the training set and *P*_*i*_ and *O*_*i*_ denotes the values for the *i*th predicted and *i*th observed phenotypes.

The Standard Deviation (SD) is a measure of how spread out numbers are.

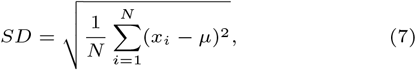

where *x* is a value in the data set, *µ* is the mean of the data set, and *N* is the number of data points in the population.

The MAE loss was watched to select the best model during the training process of 150 epochs with early stopping. The model with the lowest MAE was selected to evaluate the model on the test set as shown in Table 3 and Table 4. The SD of all the MAE values on the test set was also calculated to see the deviation among those values. The MAE measures the average absolute error over the dataset while the SD measures how far the absolute error on each training point is from the MAE. A low SD means that errors across the dataset tend to have similar values close to the mean. A high SD tells that the errors are spread over a bigger range. This can provide insight into the model: a model with a low MAE indicates a good “average” performance over the dataset, and if that model also has a low SD then it tells that the performance is not only good on average, but also uniformly on the dataset. The low values of MAE and SD of MAE for the SV and TE markers show higher prediction accuracy of `NovGMDeep` model over the SNPs.

**Table 3.**
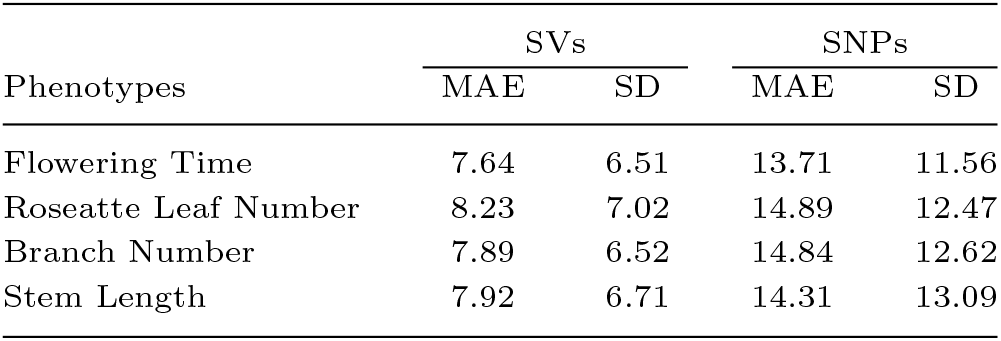
Mean Absolute Error (MAE) and Standard Deviation of MAE calculated by `NovGMDeep` model on the test sets for predicting the flowering time of *A. thaliana* using SV and SNP dataset.

| Phenotypes | SVs |  | SNPs |  |
| --- | --- | --- | --- | --- |
|  | MAE | SD | MAE | SD |
| Flowering Time | 7.64 | 6.51 | 13.71 | 11.56 |
| Rosette Leaf Number | 8.23 | 7.02 | 14.89 | 12.47 |
| Branch Number | 7.89 | 6.52 | 14.84 | 12.62 |
| Stem Length | 7.92 | 6.71 | 14.31 | 13.09 |

**Table 4.** Mean Absolute Error (MAE) and Standard Deviation of MAE calculated by `NovGMDeep` model on the test sets for predicting the flowering time of *O. sativa* using TE and SNP dataset.

| Phenotypes | TEs |  | SNPs |  |
| --- | --- | --- | --- | --- |
|  | MAE | SD | MAE | SD |
| Days to Heading | 8.65 | 6.41 | 16.84 | 18.51 |
| Plant Height | 8.01 | 6.57 | 15.92 | 19.74 |
| Spikelet Number | 9.26 | 7.22 | 19.46 | 20.25 |
| Panicle Length | 9.51 | 6.38 | 16.78 | 19.02 |

Figure 4 and Figure 5 demonstrates a typical training process for all the datasets. The trend shows that the model keeps learning the discriminative features of the data for the first few epochs as both the training and validation losses are decreasing to a significant amount at the start. For the rest of the epochs, the flattened part of the graph illustrates that the model is capable to handle the co-variate shift between the training and validation set because now the training and validation losses are decreasing steadily. The closeness of the training and validation curves demonstrates that the model is neither overfitted nor underfitted in Figure 4(a) and Figure 5(a). In Figure 4(b) and Figure 5(b) the training curve is lower than the validation curve, which shows that the model is overfitted. An underfitted model will have a high training and high validation loss while an overfitted model will have an extremely low training loss but a high validation loss. A large number of features (SNPs) expands the hypothesis space, making the data more sparse and this might also lead to overfitting problems.

**Fig. 4.**
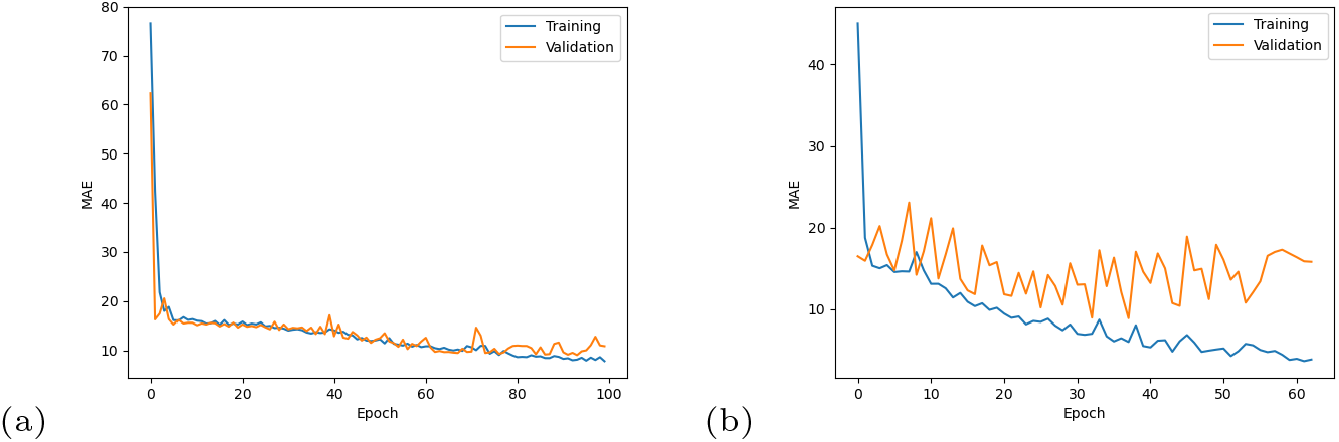
Training and validation losses of `NovGMDeep` model using (a) SV and (b) SNP dataset for predicting flowering time of *A. thaliana*. The first few epochs illustrate that the model is learning quickly as both the training and validation losses are decreasing faster. Later it shows the potential of the model in handling the co-variate shift between sets as the training and validation losses are decreasing steadily now.

**Fig. 5.**
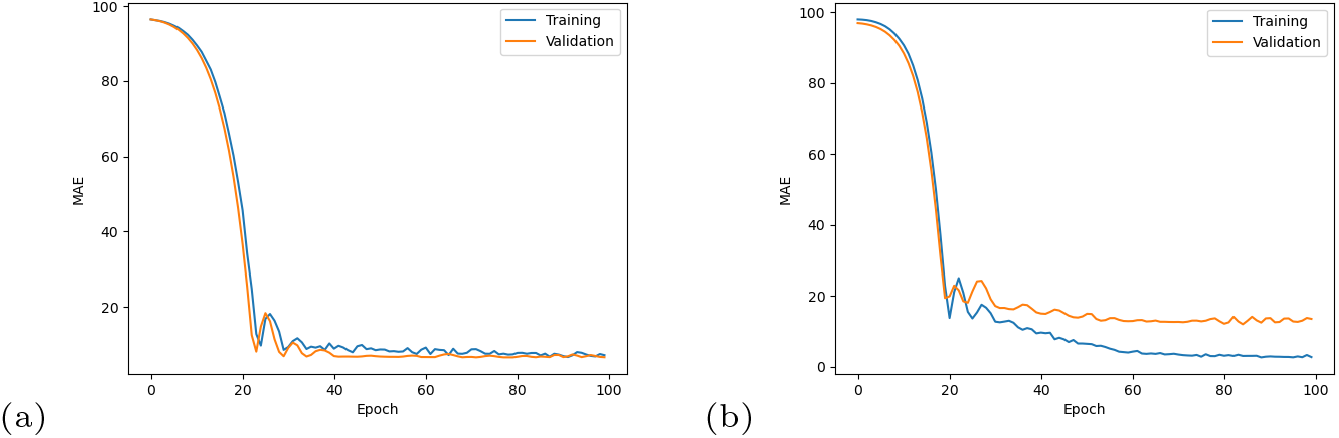
Training and validation losses of `NovGMDeep` model using (a) TE and (b) SNP dataset for predicting days to heading of *O. sativa*. The graph shows that the model is learning slowly at the start as it took 20 epochs for it to be at a steady state. Later it shows the potential of the model in handling the co-variate shift between sets as the training and validation losses are decreasing steadily now.

The `NovGMDeep` model was also compared with the aforementioned DL models in. However, the comparison was possible just on the SNP data as these models are specially designed for that. We tried the G2PDeep, which is a web-based framework where users can upload their datasets and predicts phenotypes by creating a deep learning model according to the data. Because of the extremely high-dimensional SNP dataset we used in this paper, the framework did not work well for us and, therefore, we ran the code uploaded into the repository that is publicly available in GitHub. DeepGS package was also run on the command line. PCC values were calculated for all the phenotypes of *A. thaliana* and *O. sativa* in Table 5 and Table 6 respectively.

**Table 5.** Pearson’s coefficients of correlations between the observed and predicted phenotypes on the testing set. SNPs are used to capture the genomic variations among the accessions of *A. thaliana*. Results show the average of the three PCC values taken from the three-fold cross-validation results of the models.

| Model | Flowering Time | Rosette Leaf Number | Branch Number | Stem Length |
| --- | --- | --- | --- | --- |
| *DeepGS* | 0.542 | 0.559 | 0.544 | 0.563 |
| *G2PDeep* | 0.561 | 0.557 | 0.571 | 0.569 |
| *NovGMDeep* | **0.594** | **0.563** | **0.581** | **0.572** |

**Table 6.** Pearson’s coefficients of correlations between the observed and predicted phenotypes on the testing set. SNPs are used to capture the genomic variations among the accessions of *O. sativa*. Results show the average of the three PCC values taken from the three-fold cross-validation results of the models.

| Model | Days to Heading | Plant Height | Spikelet Number | Panicle Length |
| --- | --- | --- | --- | --- |
| *DeepGS* | -0.004 | -0.015 | -0.171 | -0.165 |
| *G2PDeep* | 0.023 | -0.011 | -0.036 | -0.182 |
| *NovGMDeep* | **-0.014** | **0.002** | **-0.169** | **-0.148** |

The results in the Table 5 showed that `NovGMDeep` performed better than the other existing DL models. Also, it can be seen that G2PDeep has a better performance than DeepGS. However, in Table 6 the results for DL models suggest that statistical models are better for datasets with fewer samples as seen in Table 2.

## Discussions and Conclusions

Genomic selection has currently brought a revolution in applications of breeding programs in plants and livestock [14]. Novel prediction algorithms and methods for predicting complex traits of large genotypic data have increasingly become an essential need of breeders. High-performing deep learning techniques have served the purpose of the principal need for breeders to improve their practices. DL as an advanced technique of machine learning algorithms, has the potential to find hidden patterns in huge datasets. This work aimed to develop a DL model and evaluate its accuracy in predicting the specific traits from genome-wide SV and TE markers.

In this paper, the applications of deep learning and its relationship with GS have been explored. Although the literature suggests DL algorithms as a solid method for predicting complex phenotypes, there are a few limitations to the usage of deep learning techniques. In the existence of large-scale datasets, the most important consideration for high prediction accuracy is the model’s architecture which requires great knowledge of deep learning techniques. A well-constructed network in any model is the key source for the high performance of the model. Another crucial consideration is the choice of applying convolutional, fully connected, dropout, and sampling layers with distinct sets of hyperparameters that handle distinct characteristics of the data [4]. Interpreting the biological relevance of the data is also a helpful consideration to minimize the limitations of deep learning techniques in bioinformatics. The small number of DL for GS applications demonstrates the enormous potential for these models to enhance early candidate genotype selection and enhance knowledge of the intricate biological mechanisms underlying the link between genotypes and phenotypes. This potential is partially explained by the way such models are constructed, which provides them the capacity to recognize more intricate data patterns.

In this study, we developed `NovGMDeep` as a method to predict phenotypes based on different types of SVs (inversions, duplications, and deletions) and TEs. This work was motivated by the importance of SVs in genome evolution and that SVs could better capture variants among genomes than SNPs [30]. With the proposed model, it is still difficult to link phenotypes to genetic structural variation and environmental variables. Also, the authors of [35] concluded through GWAS that TE markers have a similar ability to discover association patterns to SNP markers. Therefore, the model is not only for predicting phenotypes from SVs and TEs but also shows that SVs can be more beneficial in genomic selection than SNPs.

Both statistical models (rrBLUP, gBLUP) have the lowest PCC values compared to the `NovGMDeep` model for the *A. thaliana* data. This is because simple regressions cannot capture the complex relationships between genotypes and phenotypes. For the *O. sativa* data the PCC value is very low for TEs and even negative for SNPs because of the low sample size to train the DL model. The results for the TE markers are better than the SNPs which makes the former more useful. Different model architectures impact the results significantly. Finding a dataset with enough samples and related phenotypic labels to train a neural network to be efficient and broadly applicable is the primary challenge that must be overcome to employ deep learning techniques for phenotype prediction. There are several datasets with significant numbers of sequenced genomes that are available, but the majority of these databases lack reliable phenotypic data for the linked samples. The cost of sequencing sufficient depth of coverage for SV and TE calls is also high. The search for datasets with more than 176 samples and associated phenotypes for TEs was unsuccessful as of the time of writing this paper. It was challenging to develop a generalizable prediction model for TEs due to the issues with overfitting and short validation sets caused by the use of datasets with a small sample size. Due to a large number of SNP markers across the genome used as features, overfitting increases in the model. This is because the number of SNPs is significantly larger than the number of samples, causing the “small-*n*-large-*p* problem”. The performance comparison between `NovGMDeep` and other DL models, such as: DeepGS [20] and G2PDeep [39], was only possible with SNP data. `NovGMDeep` outperformed the other two DL models for *A. thaliana* whereas the low sample size problem persisted for *O. sativa*.

The study also shows that the nature of data, network architecture, and the type of the trait which is being predicted play an influential role in model training and prediction. It is important to focus on the individuals with high phenotypic trait values so that they can serve as a selected asset for the different breeding programs for other vital crops. In future work, we will explore finding top-ranked individuals with high phenotypic values by incorporating all types of genomic data, such as SVs, TEs, and SNPs together. Moreover, environmental data, such as weather conditions, can be taken into consideration in phenotype prediction. Overall, the `NovGMDeep` model was well implemented in the prediction of phenotypes using SVs genotypic markers and could be added to the toolkits of crop breeders. Also, the results showed the importance of the selection of algorithms and hyperparameters for genotype-to-phenotype predictions. In conclusion, this study paves the way for the novel use of SVs and TEs in the field of GS.

## Data Sources

The phenotypes for *A. thaliana* were downloaded from https://arapheno.1001genomes.org/study/12/ and https://arapheno.1001genomes.org/study/38/.

The TE markers for *O. sativa* were downloaded from https://data.cyverse.org/dav-anon/iplant/home/yanhaidong1991/opt_genos_fltmissing_fltmaf_fltukn_TE.txt and the SNPs were downloaded from https://data.cyverse.org/dav-anon/iplant/home/yanhaidong1991/176GATK.indelsnpsFiltered0.05M0.25.txt.

The G2PDeep web-based framework can be found at https://g2pdeep.org.

The standalone G2PDeep and DeepGS packages can be found at https://github.com/kateyliu/DL_gwas and https://github.com/cma2015/DeepGSrespectively.

## Appendix: Detailed Analysis of the assumptions of PCC

Five assumptions must be met before we calculate the PCC between two variables [25]. As far as our datasets are concerned, all of the conditions are true. First, while measuring the two variables, we should consider the interval or ratio level. All the phenotypes used in the paper can describe by using intervals for example, plant height, branch number, stem length etc. on the real number line. These all are physical measures which fall into general continuous category.

Second, there must be a linear relationship between both variables. All the phenotypes when plotted using scatter plot, fell roughly along a straight line, which can also be seen in Figure 2 and Figure 3. The main thing to consider here is that the variables should not exhibit some other type of relationship, like quadratic.

Third, both variables should have a roughly normal distribution. We have plotted the values of flowering time for *A. thaliana* and days to heading for *O. sativa*. Figure 6 and Figure 7 show a bell-shaped curve for both of these phenotypes that indicates that the variables follow a normal distribution.

**Fig. 6.**
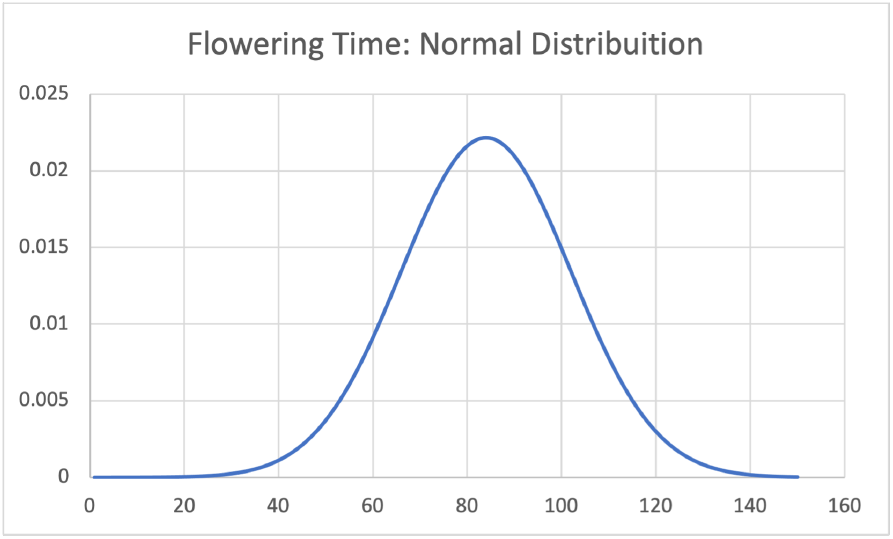
The bell-shaped curve show the values of flowering time are normally distributed.

**Fig. 7.**
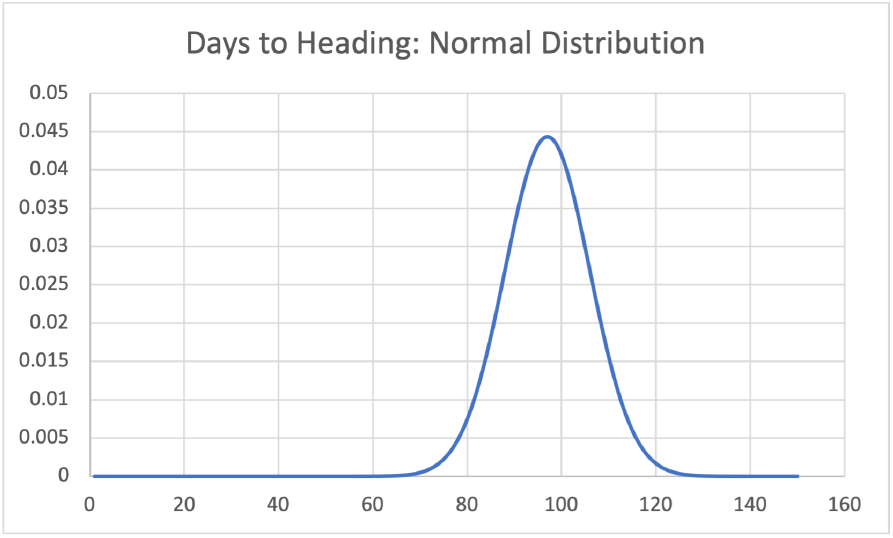
The bell-shaped curve show the values of days to heading are normally distributed.

Fourth, each observation in the dataset needs to have a pair of related values. It simply means that to calculate the correlation between the variables, each observation in the dataset has one measurement for the observed value and one measurement for the predicted value. This one was easy to check as `NovGMDeep` predicted phenotypes for all the corresponding ground truth values.

Fifth, the dataset should not contain any extreme outliers. An extreme value in the dataset substantially changes the PCC between the two variables. We did not find any extreme outlier in the dataset of predicted and observed phenotypes.

